# SpacerScope: Binary-vectorized, genome-wide off-target profiling for RNA-guided nucleases without prior candidate-site bias

**DOI:** 10.64898/2026.03.28.715005

**Authors:** Yanji Qu, Yaxuan Wang, Yan Wang, Haoru Tang, Qing Chen

**Affiliations:** College of Horticulture, Sichuan Agricultural University, Chengdu 611130, China; Key Laboratory of Agricultural Bioinformatics, Ministry of Education, Sichuan Agricultural University, Chengdu 611130, China

**Keywords:** off-target profiling, genome editing, RNA-guided nuclease, binary vectorization, indel-aware alignment

## Abstract

The precision of CRISPR/Cas systems is fundamental to their application in plant and animal biotechnology. However, comprehensive off-target assessment remains a bottleneck, particularly in large, complex genomes where existing tools often suffer from prohibitive computational costs, poor search-space convergence, and limited sensitivity toward non-canonical alignments involving insertions and deletions (indels). To address these limitations, we developed SpacerScope, an off-target analysis framework that enables unbiased, genome-wide discovery by leveraging binary vectorization and high-speed bitwise operations. Benchmarking against CIRCLE-seq data demonstrated that SpacerScope recovered 100% of validated off-target sites (6,142/6,142) within our defined parameter space, achieving zero false negatives while identifying additional high-risk sites with complex edit distances. Unlike conventional tools such as Cas-OFFinder, CHOPCHOP, and CRISPOR, SpacerScope maintains high sensitivity for indel-inclusive off-targets that are otherwise overlooked. Our results establish SpacerScope as a high-speed, high-sensitivity solution for ensuring genome-editing specificity across diverse and complex genomic landscapes. The full source code, documentation, and multi-platform executables are available at https://github.com/charlesqu666/SpacerScope.

**Graphical TOC:** SpacerScope is a genome-wide off-target profiling tool for RNA-guided nucleases that combines binary vectorization, bitwise prefiltering, compact 2-bit substitution search, and right-end-anchored indel validation. In the evaluated datasets, it recovered all 6,142 parameter-defined true off-target sites from CIRCLE-seq and showed improved detection of indel-containing candidate sites relative to the previous tools tested here. SpacerScope provides interpretable event-level annotations and empirical risk scores for downstream guide assessment.

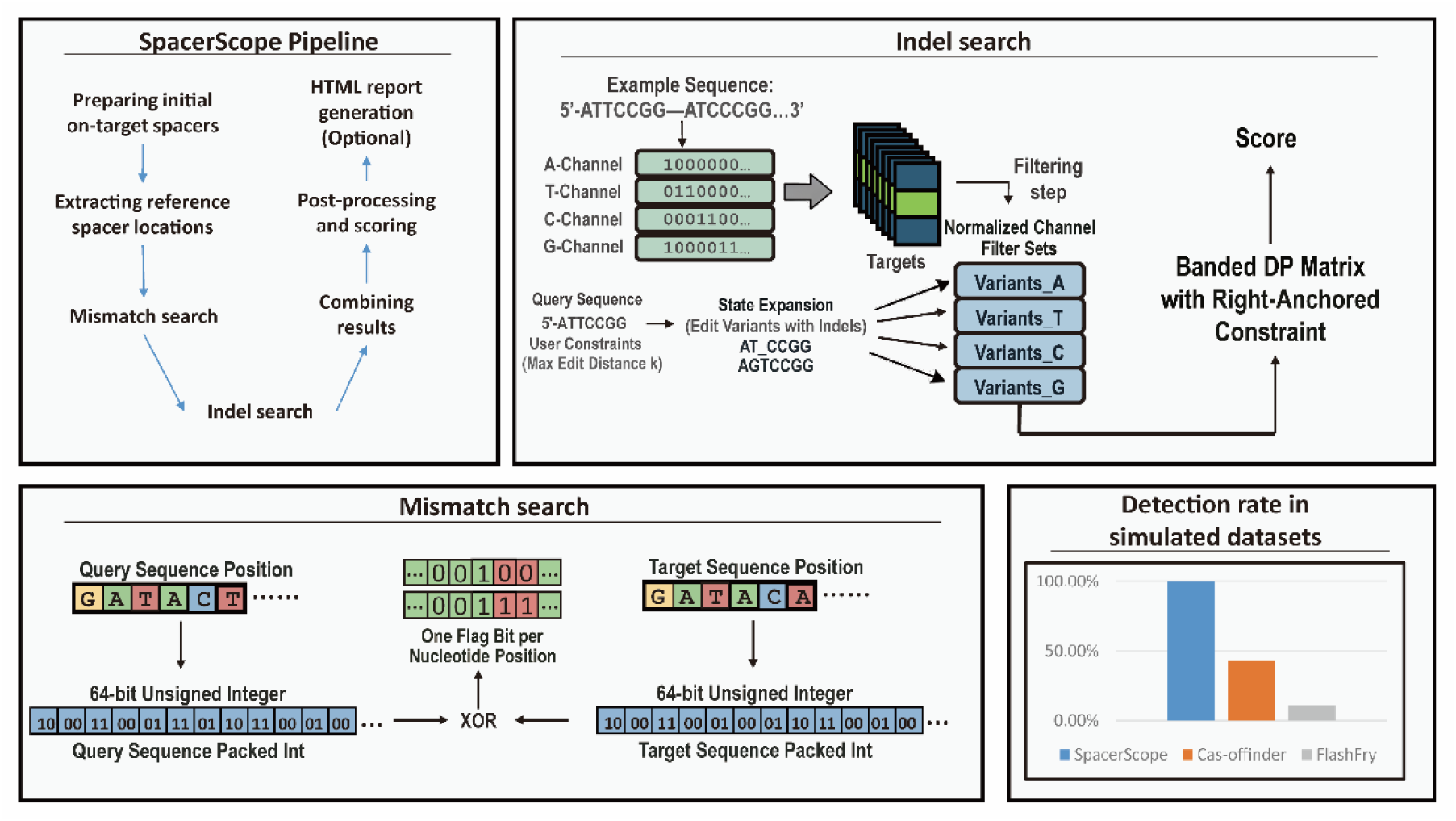

**Practitioner Points:**

> SpacerScope enables genome-wide off-target profiling with explicit support for substitutions, insertions, and deletions in a single workflow.
> The method combines binary-channel prefiltering with right-end-anchored validation to reduce the search space while preserving sensitivity within the evaluated parameter ranges.
> SpacerScope returns event-level match annotations and empirical risk scores, supporting more interpretable downstream guide selection and benchmarking.

## 1 Introduction

The diversification of Type II CRISPR/Cas systems has transformed genome engineering into a cornerstone of modern biotechnology. However, as the repertoire of CRISPR effectors and targeting rules expands, the demand for computational frameworks capable of high-precision target selection has grown increasingly acute [1, 2]. While a variety of off-target search engines have been developed, their performance often falters when faced with the inherent complexities of large-scale genomic data.

Existing algorithmic strategies generally fall into three categories, each with distinct trade-offs. Methods based on the Burrows-Wheeler Transform (BWT) or FM-index, such as Bowtie, offer efficient exact matching through compressed indexing but suffer from a precipitous decline in performance as sequence variation, particularly insertions and deletions (indels), increases [3]. Conversely, hashing and prefix-based indexing strategies, exemplified by FlashFry, facilitate rapid retrieval but can become memory-intensive on complex datasets [4]. To maximize sensitivity, exhaustive search tools like Cas-OFFinder utilize GPU acceleration via the OpenCL framework [1], yet these remain computationally expensive. Despite these advances, applying such tools to complex crop genomes remains a formidable challenge. In polyploid species characterized by high repeat content, search spaces often fail to converge, leading to excessive computational overhead and, critically, a persistent risk of undetected off-target mutations. Indeed, recent evidence suggests that many single-nucleotide variants (SNVs) induced by base editors remain beyond the predictive reach of standard tools like Cas-OFFinder [5], posing significant safety concerns for crop engineering and therapeutic applications.

A further complication arises from the inherent bias in current off-target validation workflows. Most plant studies rely on “biased” analysis—evaluating only a small subset of sites pre-selected by popular web platforms like CHOPCHOP [2] or CRISPOR [6].

This reliance on pre-filtered candidates systematically overlooks “true” off-target events residing in unflagged genomic regions, such as the highly similar paralogous gene families common in polyploid crops [7]. Furthermore, the lack of consistency in sgRNA ranking across these platforms suggests that current predictive models are not yet robust enough for standalone use [8].

To overcome these bottlenecks, we present SpacerScope, a high-performance computational framework designed to enhance the speed and fidelity of off-target analysis in complex eukaryotic genomes. SpacerScope integrates binary vectorization with a bitwise-operation-based pre-filtering mechanism, effectively eliminating the need for biased candidate-site reduction. By employing 2-bit encoding and a right-end-anchored edit distance algorithm, SpacerScope achieves exhaustive genome-wide search capabilities with significantly reduced computational latency. Our benchmarking demonstrates that SpacerScope maintains maximum sensitivity without the loss of potential target sites common in heuristic filtering. Notably, the tool excels in resolving complex edit distances where substitutions and indels coexist, ensuring more accurate mismatch quantification than conventional methods. SpacerScope is written in c++, configured to be cross-platform compatible (Windows, Linux, and macOS). It also features a web-based interface for streamlined visualization, providing a versatile and robust solution for precision genome editing across diverse biological systems.

## 2 Methods

### 2.1 Binary-Channel Filtering

To bypass the prohibitive computational overhead of exhaustive character-wise alignment, SpacerScope employs a rapid-rejection heuristic that eliminates the vast majority of non-homologous candidates before exact alignment. This binary-channel filtering strategy deconstructs sequence-matching into a high-throughput bit-pattern decision problem.

#### Encoding Architecture

Traditional DNA representation relies on four-character strings (A, T, C, G). In contrast, SpacerScope deconstructs each sequence into four discrete binary channels, one for each nucleotide. For a sequence of length ***L***, a position containing the target nucleotide is represented as ***1*** in its respective channel and ***0*** in the remaining three. For example, the sequence ‘ATCGATCG’ is encoded as an A-channel bitset of ‘10001000’, with other channels populated analogously. By transforming character-based matching into integer comparisons and bitwise operations, the algorithm leverages low-level hardware optimizations to accelerate search speed.

#### Indel-Aware Variant Filtering

To accommodate insertions and deletions (indels), SpacerScope constructs a variant filter for each query sequence within a user-defined edit distance. The algorithm systematically expands the query sequence to include all permissible substitutions, insertions, and deletions, generating a comprehensive set of “permissible variants”. To ensure uniform comparison across the reference genome, these variants, regardless of their actual length, are mapped into a fixed-length representation space. Shorter variants are normalized to maintain consistency with the canonical candidate-site length. This process yields four filter sets representing the permissible encoded values for each nucleotide channel.

#### Heuristic Rejection and Computational Efficiency

During the scanning of the reference genome, pre-encoded candidate sites are first subjected to a membership test against these filter sets. A candidate is retained for downstream exact alignment only if its encoded values in all four channels satisfy the filter conditions simultaneously. Failure in any single channel results in immediate rejection. This multi-channel pre-filtering drastically narrows the search space, ensuring that only high-probability hits proceed to computationally intensive dynamic-programming alignment. By replacing character-wise comparisons with integer-set membership tests, SpacerScope achieves high throughput and stable candidate compression, forming the core of its indel-search optimization.

#### Standardization and Handling of Non-canonical Bases

SpacerScope is optimized for standard genomic bases (A, C, G, T). During the pattern-matching phase, “N” in the Protospacer Adjacent Motif (PAM) is treated as a degenerate base. For non-standard nucleotides in the input genome or query sequences, the system treats them as “A” during 2-bit encoding but excludes them from binary-channel encoding. For transparency, these bases are restored to their original “N” designation in the final output for user verification.

To maintain optimal predictive accuracy, we recommend standard quality-control preprocessing of genomic inputs prior to analysis.

### 2.2 Right-End-Anchored Edit Distance

CRISPR/Cas-mediated cleavage is directionally constrained by the PAM. Because targeting specificity is typically highest near the PAM-proximal end, SpacerScope employs a right-end-anchored edit distance to quantify the divergence between a query sequence (***Q***) and a candidate target (***T***). While we use the term “right-end-anchored”, the algorithm dynamically adjusts orientation based on the PAM’s position relative to the spacer (5’ or 3’), which are defined by the users. This metric functions as a semi-global extension of the classical Levenshtein distance. Unlike standard global alignment, our approach permits a free offset at the PAM-distal (left) end. Prefixes that do not contribute to the optimal alignment are excluded from the edit cost, while the PAM-proximal (right) end remains strictly anchored. This asymmetric constraint effectively models length variations induced by indels while maintaining alignment fidelity in the biologically critical region adjacent to the PAM. By allowing sequences to initiate alignment from different starting points at the distal end, SpacerScope avoids inflating the edit distance with penalties caused by simple sequence offsets, focusing instead on true substitution and indel events.

### 2.3 Computational Workflow

#### 2.3.1 Low-Complexity Sequence Filtering

To mitigate non-specific binding and enhance computational throughput, all candidate spacers undergo a low-complexity filtration step. We implement a configurable DUST-like heuristic based on trinucleotide frequencies to identify and exclude highly repetitive genomic regions. For each trinucleotide ***t,*** a penalty score is calculated based on its occurrence ***nt***. Only sequences exceeding a user-defined complexity threshold are retained, ensuring that the resulting sgRNA candidates possess sufficient information density for unique genomic targeting. A large number can be set to disable this filtering process.

#### 2.3.2 High-Throughput Substitution Search

For search scenarios restricted to base substitutions, SpacerScope utilizes a bitwise-parallel workflow leveraging compact 2-bit encoding. Both query and target sequences are mapped to binary representations (A=00, C=01, G=10, T=11), enabling a 20–30 bp spacer to be encapsulated within a single 64-bit unsigned integer. For targets on the antisense strand, the algorithm computes the reverse complement prior to encoding to maintain a unified search orientation. Mismatch quantification is performed via an optimized bitwise logic: An XOR operation identifies bit-level discrepancies between the query and target. Discrepancies are masked and compressed into a single-bit-per-nucleotide flag format. The total mismatch count is retrieved using the hardware-accelerated population count (popcount) instruction. By bypassing character-wise traversal in favor of basic bitwise primitives (XOR, SHIFT, MASK, and POPCOUNT), SpacerScope achieves near-theoretical maximum throughput for substitution scanning.

##### Batch Processing and Memory Optimization

To further maximize execution speed, we implemented a batch-scanning architecture. Rather than iterating through the reference genome for each individual query, which would incur prohibitive I/O overhead, SpacerScope groups multiple queries into a single processing batch. This allows the target set to be scanned sequentially, significantly improving cache locality and reducing redundant memory access. Pre-encoded targets are stored in contiguous arrays to ensure streamlined sequential reads, while the core comparison loops are unrolled to minimize branch-prediction penalties. The integrated substitution-search workflow is summarized in Figure 1.

**Figure 1.**
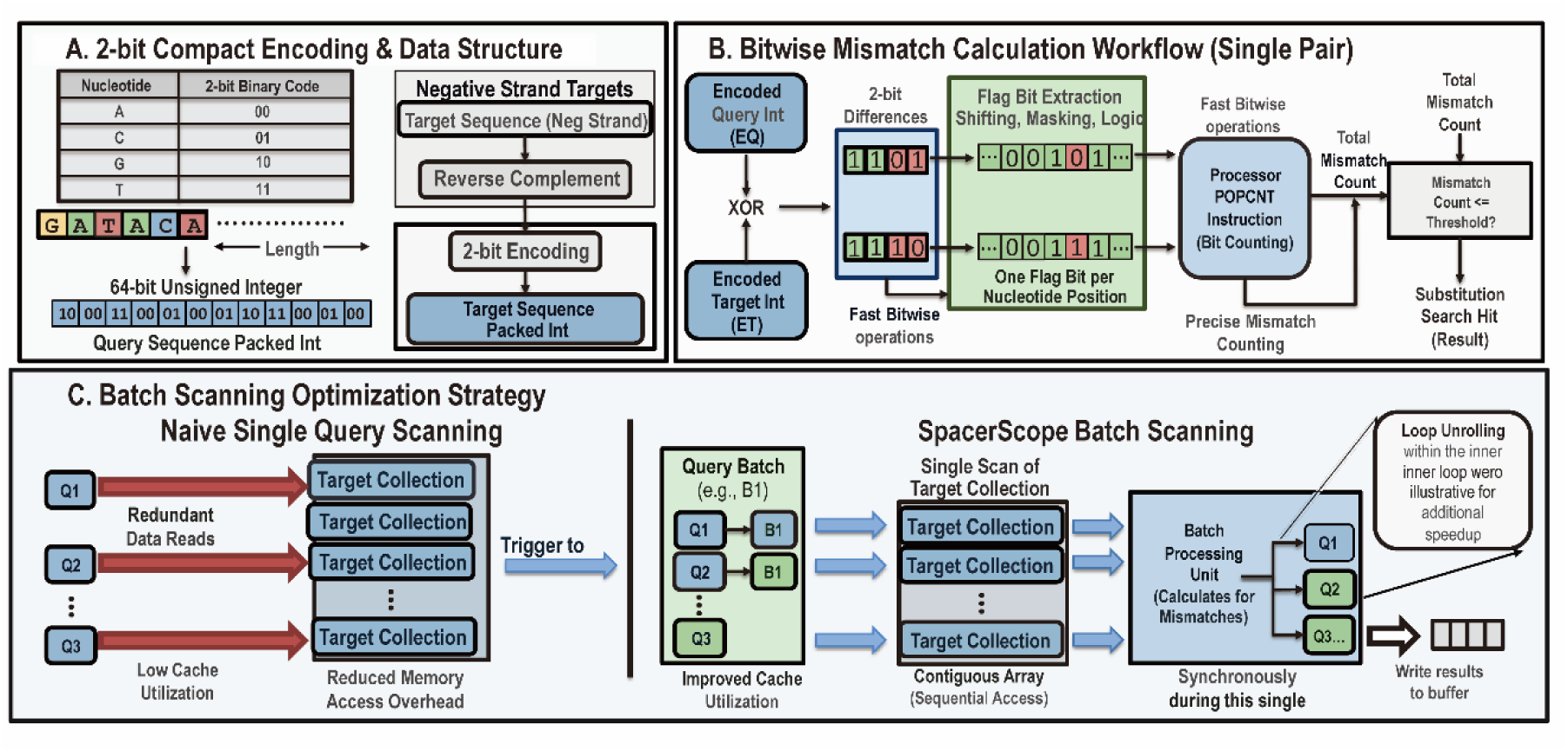
Schematic workflow of substitution search.

#### 3.2.3 Indel-Aware Search Strategy

Unlike substitution-only scanning, the inclusion of insertions and deletions (indels) precludes the use of simple bitwise XOR for mismatch quantification. To maintain high-throughput performance without compromising sensitivity, SpacerScope employs a two-stage heuristic-to-exact pipeline: (1) high-speed Binary-Channel Prefiltering to prune the search space, followed by (2) Right-End-Anchored Banded Dynamic Programming (DP) for rigorous validation. The integrated indel-search workflow is illustrated in **Figure 2**.

**Figure 2:**
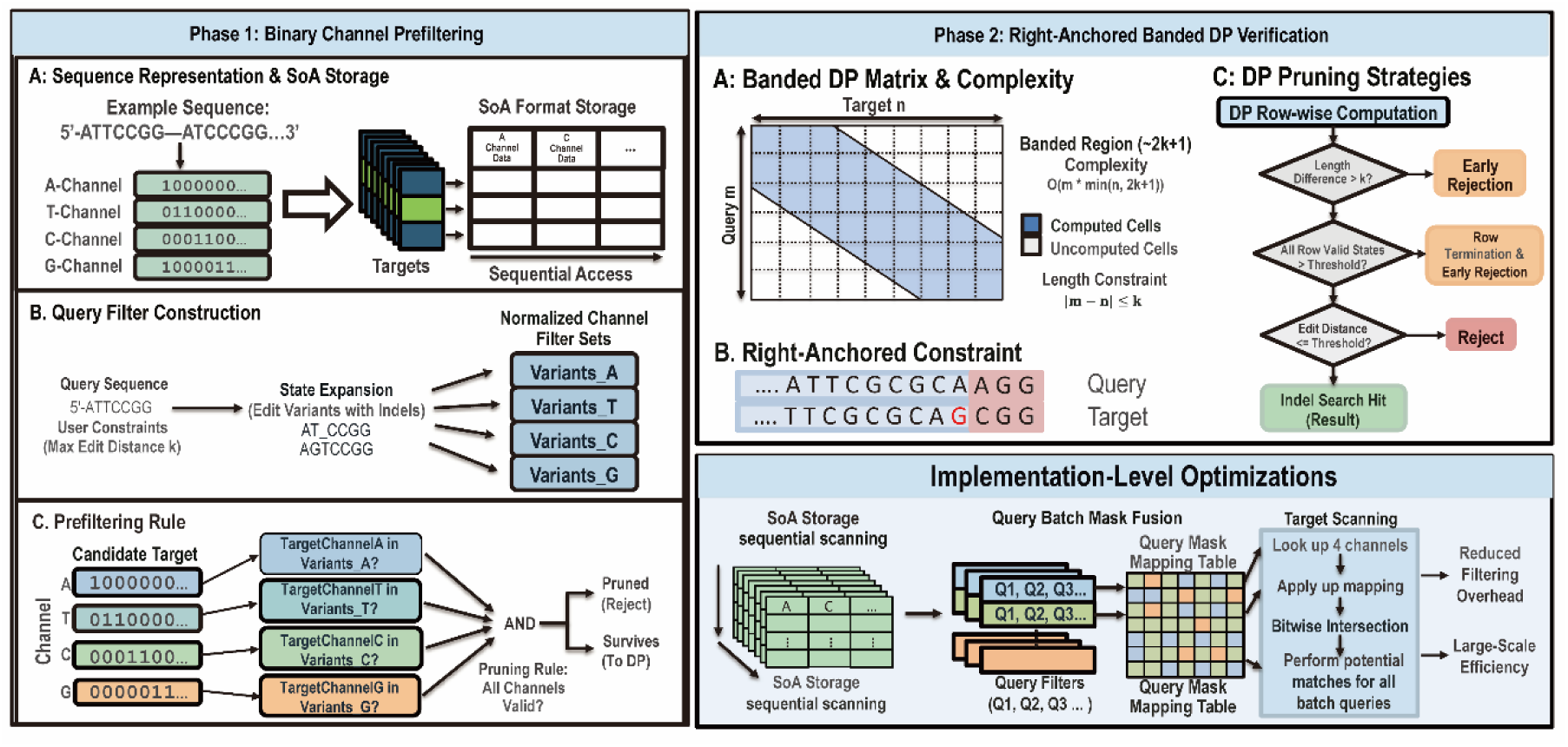
Schematic workflow of indel search.

##### Binary-Channel Prefiltering

The prefiltering stage abstracts genomic sequences into a one-hot bitset representation. Each sequence is decomposed into four discrete binary channels (A, C, G, and T). A bit value of 1 at a specific position indicates the presence of the corresponding nucleotide, while the remaining channels are set to 0. This transformation converts the traditional string-matching problem into a multi-channel pattern recognition task. For each query, SpacerScope pre-calculates the “edit reachability” within the user-defined maximum edit distance. The algorithm generates all permissible variants, accounting for substitutions and indels, and maps them into a fixed-length channel space. This results in four comprehensive filter sets: ***FA***, ***FC***, ***FG*** and ***FT***. A candidate genomic site is retained for exact alignment only if its encoded values for all four channels belong to their respective filter sets:

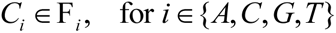

Any candidate failing this membership test is immediately pruned. This design ensures that the computationally expensive DP alignment is reserved only for high-probability candidates, effectively pre-encoding the complexity of indels into a rapidly evaluable bitmask.

##### Right-End-Anchored Banded Dynamic Programming

Candidates surviving the prefiltering stage undergo exact validation via a specialized semi-global DP algorithm. To minimize the computational footprint, we implement a banded DP approach. Given a query of length ***m***, a target of length ***n***, and a maximum edit distance ***k***, any candidate where |m - n| > k is discarded immediately. For eligible candidates, the algorithm computes only a diagonal band of width approximately 2k + 1, reducing the time complexity from the standard O(mn) to O(mk). This validation stage is uniquely constrained by right-end anchoring, prioritizing alignment stability at the PAM-proximal end. We further optimize the DP execution through two early-termination rules: Immediate rejection if the cumulative length divergence exceeds the threshold ***k***. Termination of the current alignment if all valid states within a given DP row exceed the maximum permissible edit distance. Consequently, the banded DP is executed over a highly restricted local state space, ensuring that “true” hits are validated with maximum precision and minimal latency.

##### Implementation of Low-Level Optimizations

To bridge the gap between algorithmic design and hardware performance, SpacerScope introduces several low-level optimizations. Structure of Arrays (SoA) Layout: Genomic targets are stored in a contiguous SoA format by nucleotide channel. This layout maximizes spatial locality and minimizes cache misses during sequential genome-wide scans. Batch Mask Fusion: When processing multiple queries simultaneously, individual channel filters are merged into unified lookup tables. These tables map channel values to query bitmasks, allowing the system to evaluate a batch of queries through a single set of bitwise intersections.

This hardware-aware architecture, combined with our pruning heuristics, enables SpacerScope to perform exhaustive off-target discovery at speeds comparable to heuristic-based tools.

#### 2.3.4 High-Fidelity Alignment and Empirical Risk Scoring

While the initial scanning stages identify potential query-target associations, these raw outputs represent low-resolution hits that lack the granularity required for formal off-target classification and risk assessment. SpacerScope therefore implements a dedicated post-processing module to perform high-fidelity sequence parsing, heuristic-driven scoring, and event-level interpretation.

##### Contextual Sequence Retrieval

The module first consolidates raw candidates from both substitution and indel search streams to calculate genomic occurrence frequencies. To overcome the limitations of fixed-length search windows, which can lead to truncated alignments in the presence of indels, the post-processing engine retrieves the full genomic context, including flanking regions, directly from the reference assembly. By reparsing chromosome coordinates and strand orientation from the original headers, SpacerScope ensures that the subsequent alignment is conducted within a biologically complete sequence landscape.

##### Constrained Event-Enumeration

To ensure biological relevance, SpacerScope utilizes a constrained event-enumeration procedure. Rather than relying on a singular, generic local alignment, the algorithm recursively explores all permissible interpretation paths (substitutions, insertions, and deletions) within the user-defined edit-distance threshold. This exhaustive traversal records the precise coordinates and base-pair identities of every editing event. This approach ensures that the resulting alignment is not merely a mathematical optimum, but a biologically plausible interaction between the sgRNA and its genomic target.

##### Empirical Risk Scoring

Candidate hits are prioritized using a heuristic risk-scoring function calibrated from validated off-target observations compiled in [9] and subsequently refined through empirical adjustments. The score integrates three primary considerations: Event Type, reflecting the differential impact of substitutions and indels; Positional Bias, reflecting the greater sensitivity of the PAM-proximal seed region; and Event Adjacency, reflecting the compounded effect of neighboring mismatches. When multiple biologically plausible interpretation paths exist for a single site, SpacerScope retains the highest-scoring trajectory as the representative alignment. Accordingly, the reported Score and Details fields are intended to reflect the most plausible relative risk profile rather than an arbitrary alignment output. This score should therefore be interpreted as a comparative risk index, not as an experimentally calibrated absolute probability.

##### Classification and Reporting

Following alignment optimization, hits are categorized into three distinct classes: A perfect genomic match, a match containing only substitutions or a match containing at least one insertion or deletion. The final distance metric is updated to reflect the true number of divergence events identified during refined alignment. Results are exported as a structured, nine-field TSV table (including QueryName, GenomicFrequency, and Score), which serves as the data source for an automated, interactive HTML visualization report.

## 3 Results

### 3.1 Benchmarking via Simulated Genomic Datasets

To rigorously evaluate the algorithmic fidelity and sensitivity of SpacerScope across a spectrum of editing topologies, we first performed validation using two simulated 1-Mb genomic datasets. These in silico benchmarks were engineered to include a ground-truth repertoire of heterogeneous mismatch profiles, encompassing single insertions coupled with substitutions, tandem insertions, multiple substitutions, and simultaneous insertion-deletion events.

In Dataset A (low-frequency implantation), 413 ground-truth sites were embedded (approx. 1 site per 2.5 kb) to simulate a sparse off-target landscape. In Dataset B (high-density implantation), 20,019 sites were embedded (approx. 1 site per 50 bp) to assess algorithmic performance under extreme search-space saturation. Following analysis, SpacerScope successfully recovered 100% of the implanted ground-truth sites in both datasets (426 and 20,254 total hits detected, respectively), confirming the robustness of the bitwise-filtering and banded-DP pipeline across varied genomic densities (**Figure 3**).

**Figure 3:**
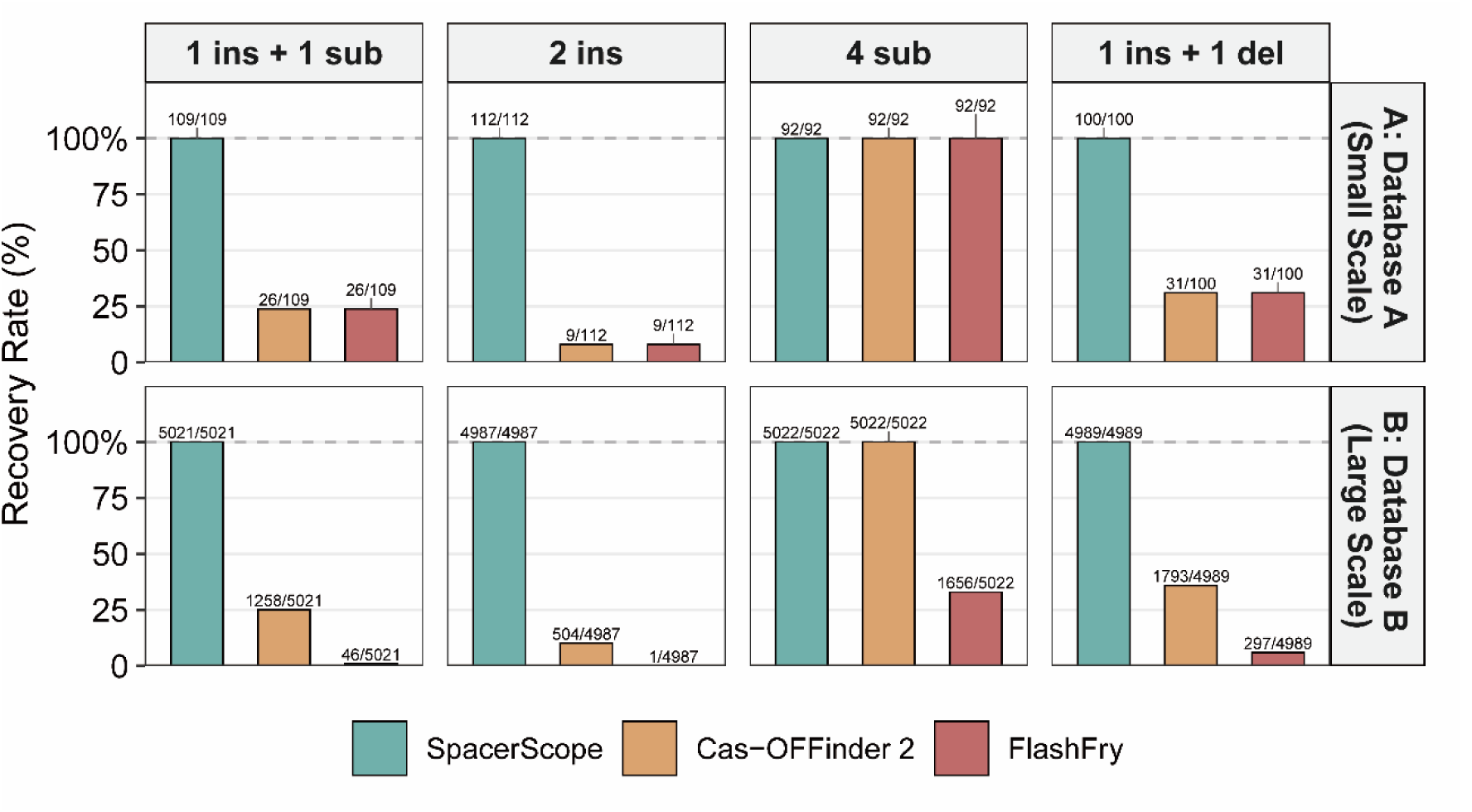
Composition of the simulated datasets and matching results used to validate implementation completeness and coverage of event types. Note: Ins, insertion; Del, deletion; Sub, substitution. All indel and substitution events are defined relative to the target sequence compared to the query spacer.

To contextualize these results, we subjected the same synthetic benchmarks to analysis by established tools, including Cas-OFFinder 2 and FlashFry. Both frameworks exhibited a significant sensitivity gap, failing to recover a substantial subset of the ground-truth sites. This performance divergence was particularly pronounced in match types involving non-canonical indel-inclusive alignments, where existing tools reached their inherent detection limits (Figure 3). These findings underscore the superior capacity of SpacerScope to resolve complex edit distances that are systematically overlooked by current industry-standard software.

### 3.2 Validation via Empirically Verified Off-Target Sites in *Oryza sativa*

To evaluate the predictive accuracy of SpacerScope in a complex crop genome, we leveraged a high-confidence off-target dataset from a previous whole-genome sequencing (WGS) study in rice (*Oryza sativa*) [10]. Tang et al. characterized the off-target profiles of several sgRNAs, identifying 12 empirically validated off-target sites for a guide targeting the Os02circ25329 locus (5’-GCAGCTCTGACATGTGGGCCCGG-3’). Using this sequence as a query, we screened the rice reference genome with SpacerScope, applying a search threshold of 4 mismatches and 2 indels. SpacerScope identified 180 candidate loci, achieving 100% sensitivity by successfully recovering all 12 previously validated off-target sites. Notably, the Unique Editing Rate (UER), a metric integrated into SpacerScope to quantify targeting precision, was calculated at a mere 0.56% for this sgRNA. This low value is highly concordant with the extensive off-target activity observed experimentally, highlighting the tool’s utility in identifying high-risk guides during the design phase.

We further extended our analysis to the remaining 11 sgRNAs described in the same study. For these guides, the original authors reported negligible off-targeting or variations attributable to background genetic noise rather than Cas9 activity. Consistent with these biological observations, SpacerScope assigned 7 of these sgRNAs a UER exceeding 90%, with the lowest recorded value at 64.76%. These results demonstrate a strong correlation between SpacerScope’s predictive metrics and in vivo editing outcomes. A comprehensive summary of the SpacerScope predictions for the rice genomic targets is provided in **Table 1**.

**Table 1.** Off-target prediction summary for rice sgRNAs.

|  | Total | Exact | Mismatch | Indel | Unique Editing Rate (UER) |
| --- | --- | --- | --- | --- | --- |
| Os03circ00204-sgRNA02 | 14 | 1 | 12 | 1 | 96.41% |
| Os03circ00204-sgRNA01 | 19 | 1 | 17 | 1 | 91.53% |
| Os02circ25329-sgRNA02 | 20 | 1 | 18 | 1 | 92.98% |
| Cas9-B: | 29 | 1 | 27 | 1 | 93.80% |
| Cas9-F | 32 | 2 | 27 | 3 | 91.97% |
| Cas9-G | 40 | 2 | 36 | 2 | 96.52% |
| Cas9-D | 53 | 2 | 45 | 6 | 73.89% |
| Cas9-E | 57 | 2 | 50 | 5 | 71.09% |
| Cas9-H | 63 | 1 | 61 | 1 | 90.91% |
| Cas9-C | 179 | 1 | 165 | 13 | 64.76% |
| Os02circ25329-sgRNA01 | 180 | 5 | 152 | 23 | 0.56% |
| Cas9-A | 187 | 1 | 184 | 2 | 78.46% |

Note: The "Unique Editing Rate (UER)" is a relative specificity measure summarized from SpacerScope empirical risk scores and is intended to quantify the relative likelihood that a candidate sgRNA edits only its perfectly matched site, rather than an experimentally calibrated absolute probability. Total denotes the total number of predicted matching sites; Exact denotes the number of perfectly matched sites; and Mismatch/Indel denote the number of sites containing mismatches or insertions/deletions. Os02circ25329-sgRNA01 is the sgRNA for which Xu Tang et al. validated 12 true off-target sites.

We also performed a comparative benchmark using Cas-OFFinder 2 and FlashFry on the rice genome. All tools were executed in single-threaded mode to ensure a fair comparison of algorithmic efficiency. Because the 12 previously validated off-target sites in rice [10] were characterized by low sequence divergence (substitution-only), all three tools successfully recovered these primary hits. However, SpacerScope consistently identified a broader range of potential off-target candidates across 11 of the 12 tested sgRNAs (**Table 2**). This expanded detection capability is largely attributed to SpacerScope’s superior handling of indel-inclusive alignments, which are frequently overlooked by conventional heuristic-based tools. While Cas-OFFinder 2 remains marginally faster due to its highly optimized substitution-only logic, its inability to detect complex indels limits its utility in high-precision applications. Conversely, FlashFry exhibited significantly higher computational latency (668s) compared to SpacerScope (64s), highlighting the efficiency of our bitwise-parallel architecture.

**Table 2.** Comparative Benchmarking of sgRNA Targeting in the Rice Genome.

|  | SpacerScope (fasta mode) | SpacerScope (fasta-cut mode) | Cas-OFFinder 2 | FlashFry |
| --- | --- | --- | --- | --- |
| Total detected sites | 873 | 873 | 832 | 832 |
| Os03circ00204-sgRNA02 | 14 | 14 | 13 | 13 |
| Os03circ00204-sgRNA01 | 19 | 19 | 18 | 18 |
| Os02circ25329-sgRNA02 | 20 | 20 | 19 | 19 |
| Cas9-B | 29 | 29 | 28 | 28 |
| Cas9-F | 32 | 32 | 29 | 29 |
| Cas9-G | 40 | 40 | 39 | 39 |
| Cas9-D | 53 | 53 | 52 | 52 |
| Cas9-E | 57 | 57 | 57 | 57 |
| Cas9-H | 63 | 63 | 62 | 62 |
| Cas9-C | 179 | 179 | 173 | 173 |
| Os02circ25329-sgRNA01 | 180 | 180 | 157 | 157 |
| Cas9-A | 187 | 187 | 185 | 185 |
| Peak memory usage (GB) | 15.6 | 13.2 | 13.7 | 14.7 |
| Runtime (s) | 64 | 63 | 47 | 668 |

Note: Performance was evaluated on a standardized environment; peak memory reflects system-wide overhead during execution. SpacerScope (FASTA-cut mode) demonstrates optimized memory efficiency without sacrificing detection sensitivity.

### 3.3 Sensitivity Benchmarking Against CIRCLE-seq Experimental Data

To assess the reliability of SpacerScope in a large, complex genomic context, we utilized the human CIRCLE-seq dataset [11], a representative in vitro benchmark dataset for off-target evaluation. We configured SpacerScope with inclusive parameters (6 mismatches, 2 indels) to target both canonical (NGG) and non-canonical (NAG) PAM sites. The results confirm the exceptional sensitivity of the SpacerScope framework. Within the defined parameter space (6,142 CIRCLE-seq validated sites), SpacerScope achieved zero false negatives, recovering 100% of true off-targets. In contrast, existing tools exhibited a severe sensitivity deficit under identical conditions (**Figure 4**). Furthermore, as the mismatch threshold increased to 6, both FlashFry and Cas-OFFinder showed a dramatic decrease in the recovery of low-mismatch variants, likely due to heuristic pruning errors or alignment failures. SpacerScope’s corrective post-processing mechanism and exhaustive pre-filtering ensure that these high-risk sites are retained, making it the most robust tool for comprehensive off-target risk assessment in human and complex crop genomes.

**Figure 4:**
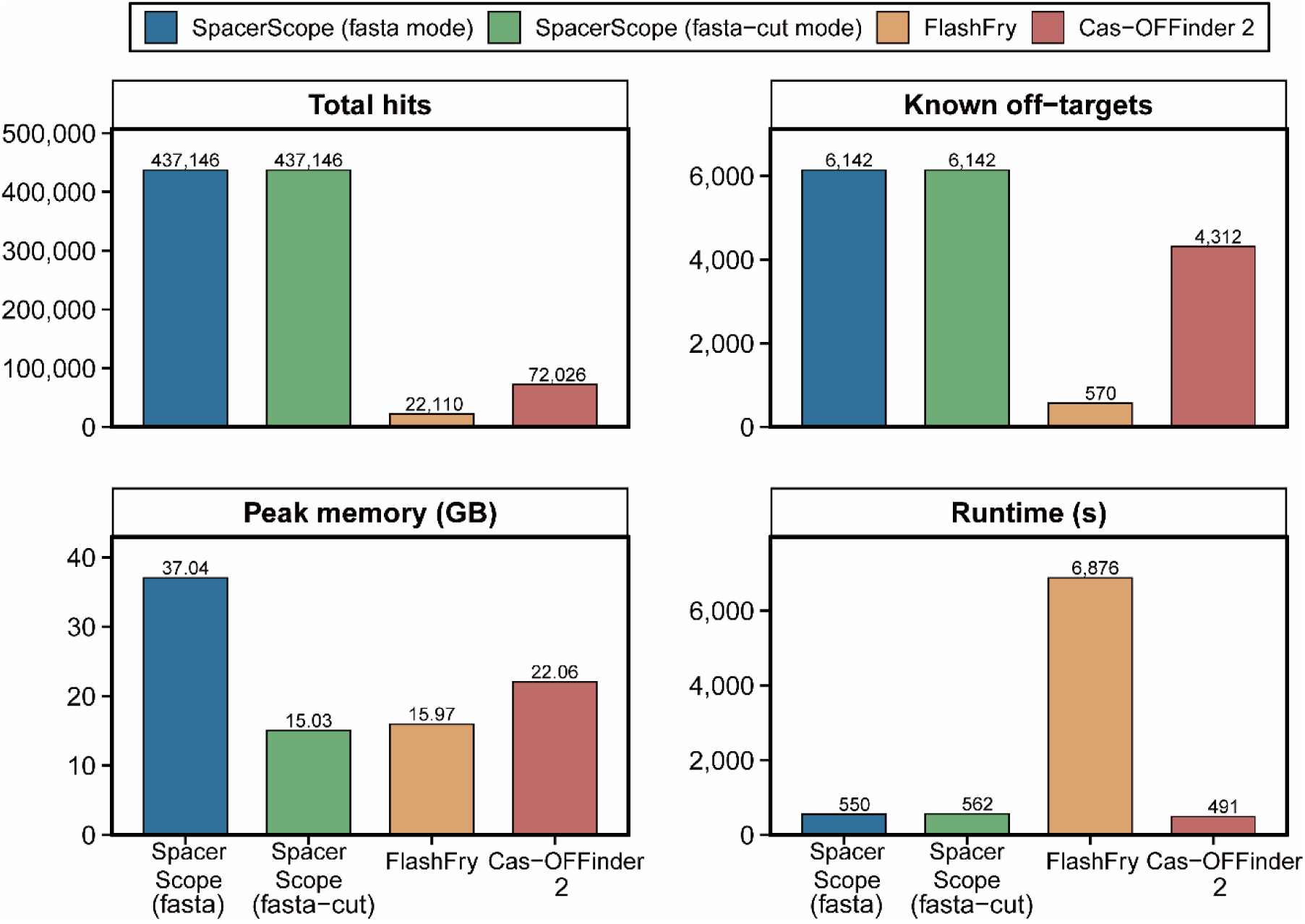
Benchmarking Against CIRCLE-seq data using SpacerScope. Note: the reported peak memory values reflect overall system-wide peak usage during runtime rather than the true memory consumption of the individual tools, and are provided only for relative comparison.

### 3.4 Comparative Analysis with Web-Based Design Platforms

To evaluate the practical utility of SpacerScope against established workflows, we conducted cross-platform comparisons with CHOPCHOP v2 [2] and CRISPOR [6]. We selected four representative loci across mammalian and plant models: EXT1 (human and mouse), AGAMOUS (Arabidopsis), and its rice ortholog OsMADS3. These genes were selected for their fundamental roles in skeletal development (EXT1) and reproductive fitness (AGAMOUS/OsMADS3), representing high-priority targets for both therapeutic and agricultural engineering [12, 13].

Using the default CHOPCHOP v2 algorithm, we identified the top-ranked sgRNA for each locus and re-evaluated their off-target profiles using SpacerScope. Our analysis revealed that SpacerScope consistently identified a greater number of potential off-target sites, particularly as sequence divergence increased (**Table 3**). For instance, in the human EXT1 locus, CHOPCHOP identified no off-targets within its maximum threshold (MM < 3), whereas SpacerScope detected four potential sites. This discrepancy highlights the sensitivity advantage of SpacerScope’s exhaustive bitwise-parallel search over the heuristic-limited indices used by general web platforms.

**Table 3.**
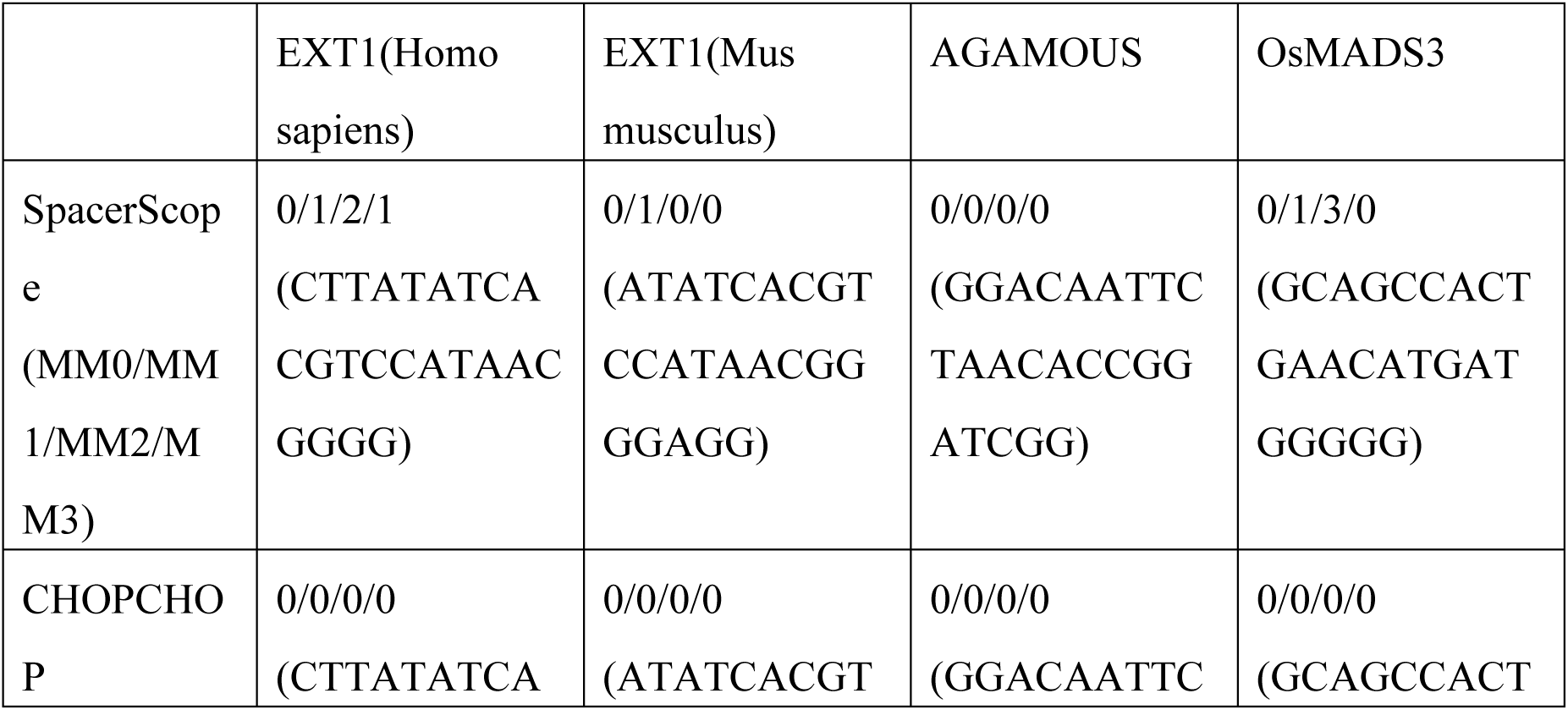
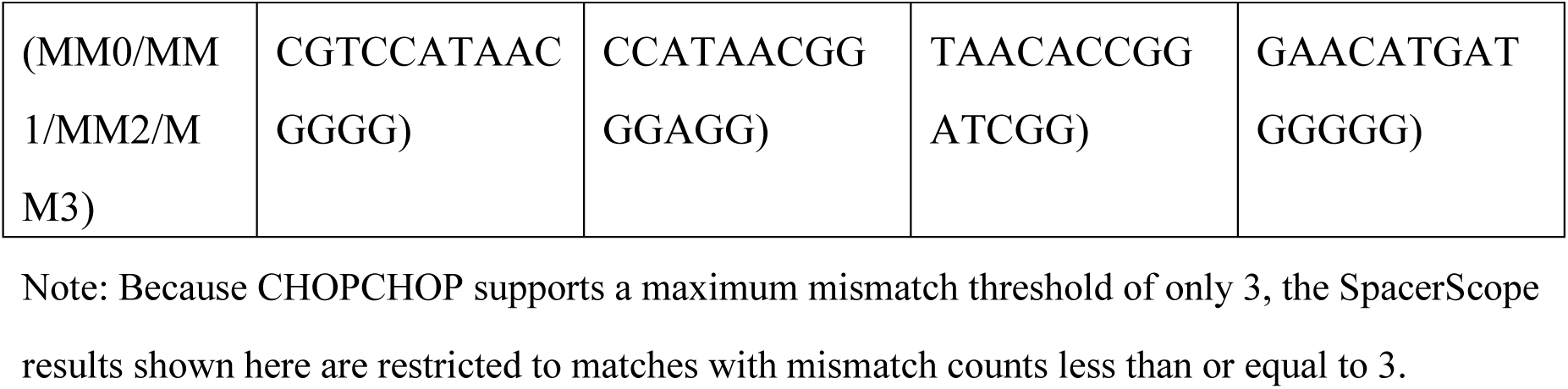
Representative gRNA case comparisons between SpacerScope and CHOPCHOP across different species.

Similar results were observed when benchmarking against CRISPOR [6]. Due to CRISPOR’s input length restrictions (< 2,300 bp), only partial gene fragments were used for comparative analysis. Despite the use of updated reference assemblies for Arabidopsis (Ler) and rice (Nipponbare), SpacerScope identified significantly more potential off-target loci, often an order of magnitude higher in the MM4 category (**Table 4**). These findings underscore that web-based tools, while accessible, may systematically underestimate off-target risks by employing restrictive candidate-site filtering.

**Table 4.**
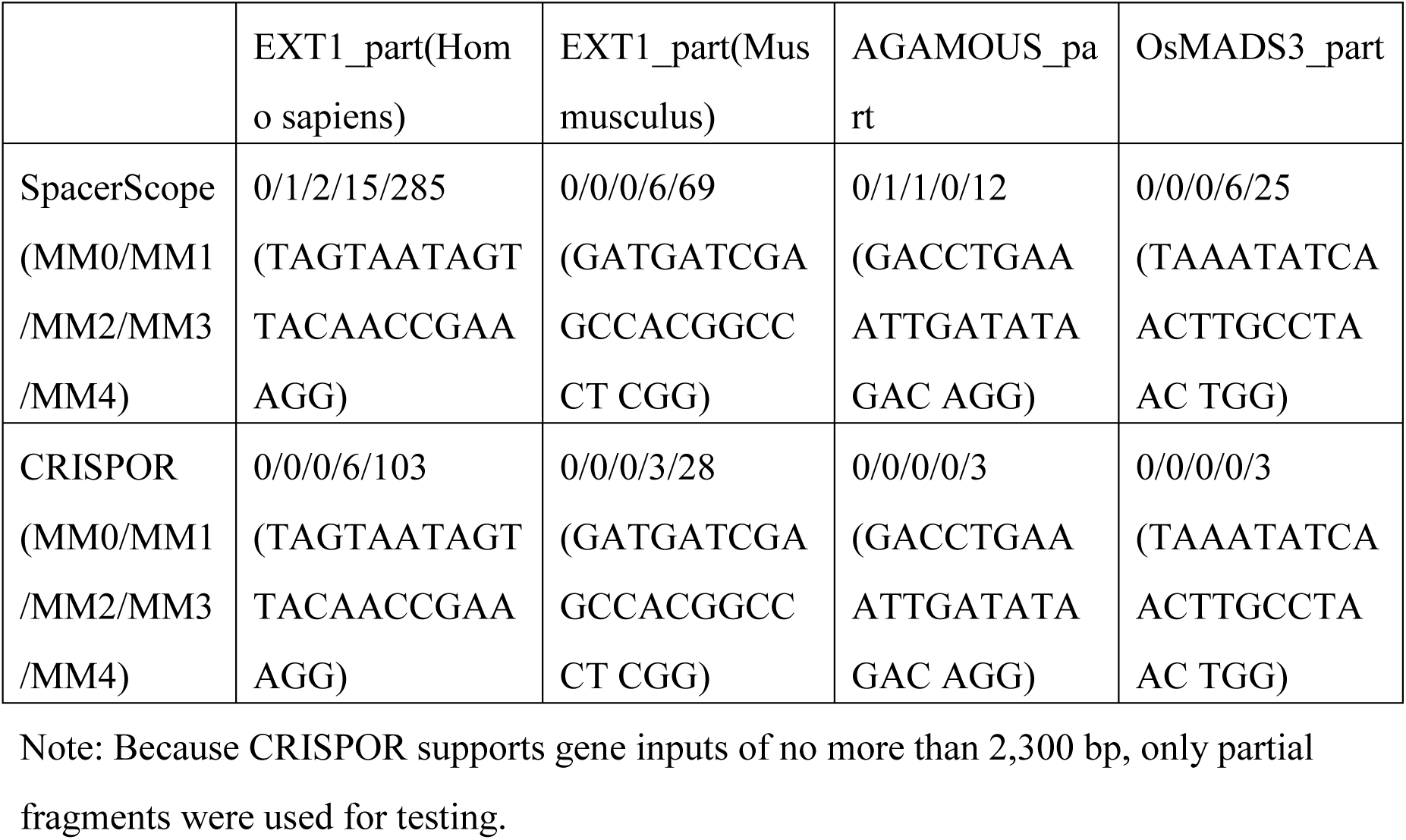
Representative gRNA case comparisons between SpacerScope and CRISPOR across different species.

### 3.5 Evaluation of Targeting Fidelity in Polyploid Genomes

Off-target prediction is particularly challenging in polyploid species due to high homeologous sequence similarity. We compared SpacerScope with CRISPR-PLANT v2 [14] using the *MSH1* gene (a regulator of organellar genomic stability) as an example in the octoploid strawberry (*Fragaria* × *ananassa* cv. Benihoppe) genome [15]. Our analysis revealed a notable divergence in target categorization. Several candidate sgRNAs assigned the highest recommendation rating (“A1”) by CRISPR-PLANT v2 were found by SpacerScope to have a UER of less than 70% (**Table 5**). This suggests that while these sites may appear optimal under standard global/local alignment parameters, SpacerScope’s exhaustive search uncovers a significant burden of potential off-targets in the complex strawberry genome. These results highlight the necessity of employing high-sensitivity tools like SpacerScope when navigating the redundant genomic landscapes of polyploid crops.

**Table 5.**
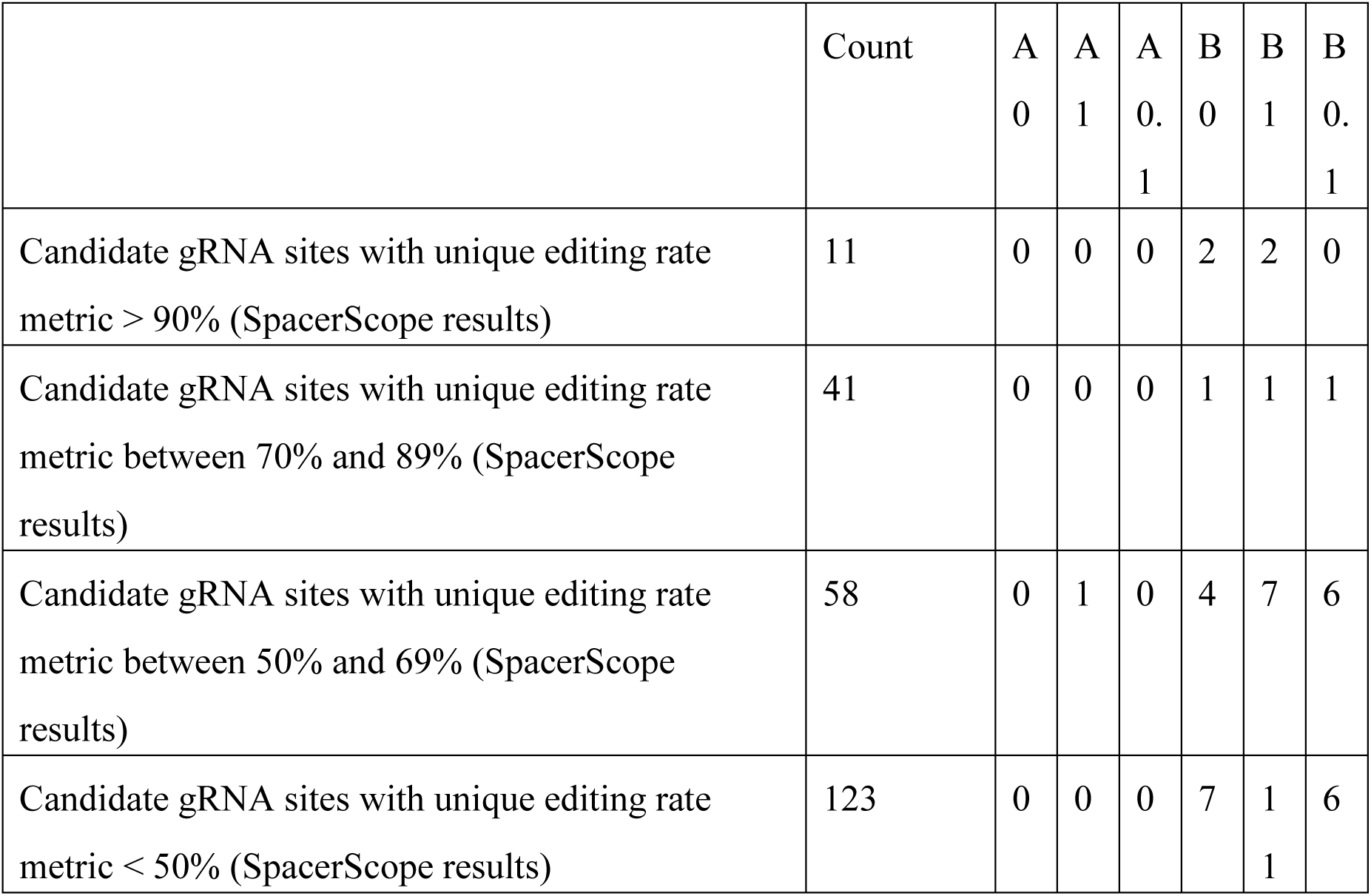
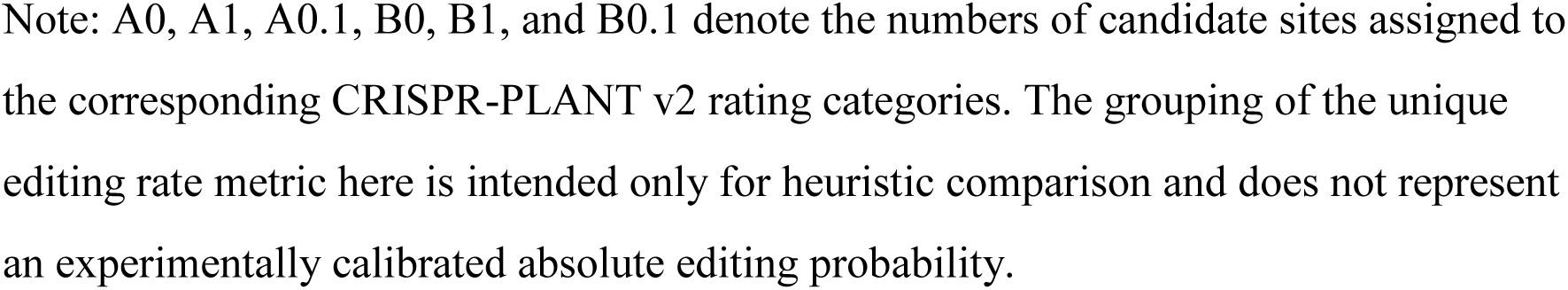
Overlap between SpacerScope msh1 matching results and CRISPR-PLANT v2 ratings in *Fragaria* × *ananassa*.

|  | Count | A<br>0 | A<br>1 | A<br>0.<br>1 | B<br>0 | B<br>1 | B<br>0.<br>1 |
| --- | --- | --- | --- | --- | --- | --- | --- |
| Candidate gRNA sites with unique editing rate metric > 90% (SpacerScope results) | 11 | 0 | 0 | 0 | 2 | 2 | 0 |
| Candidate gRNA sites with unique editing rate metric between 70% and 89% (SpacerScope results) | 41 | 0 | 0 | 0 | 1 | 1 | 1 |
| Candidate gRNA sites with unique editing rate metric between 50% and 69% (SpacerScope results) | 58 | 0 | 1 | 0 | 4 | 7 | 6 |
| Candidate gRNA sites with unique editing rate metric < 50% (SpacerScope results) | 123 | 0 | 0 | 0 | 7 | 1<br>1 | 6 |

Note: A0, A1, A0.1, B0, B1, and B0.1 denote the numbers of candidate sites assigned to the corresponding CRISPR-PLANT v2 rating categories. The grouping of the unique editing rate metric here is intended only for heuristic comparison and does not represent an experimentally calibrated absolute editing probability.

### 3.6 Computational Scaling and Performance Benchmarking

To assess the resource footprint and scalability of SpacerScope, we performed multi-threaded benchmarks using full-length AGAMOUS and EMX1 sequences against the Arabidopsis and human genomes, respectively. Benchmarking was conducted on a high-performance computing (HPC) node equipped with dual AMD EPYC 7R32 processors (96 physical cores).

SpacerScope demonstrated robust computational scaling, with runtimes decreasing significantly as thread counts increased (**Figure 5A**). In the human genome scan, SpacerScope maintained high throughput even when exhaustive indel-search was enabled, a testament to the efficiency of the bitwise pre-filtering architecture. Peak memory usage was observed to be stable and predictable across varied workloads (**Figure 5B**). While system-wide memory peaks reached approximately 13–15 GB, the baseline resident memory remained well within the capacity of modern workstations and HPC environments. These results establish SpacerScope as a high-efficiency solution capable of handling large-scale genomic data with a manageable resource footprint, making it suitable for both institutional servers and individual research workstations.

**Figure 5:**
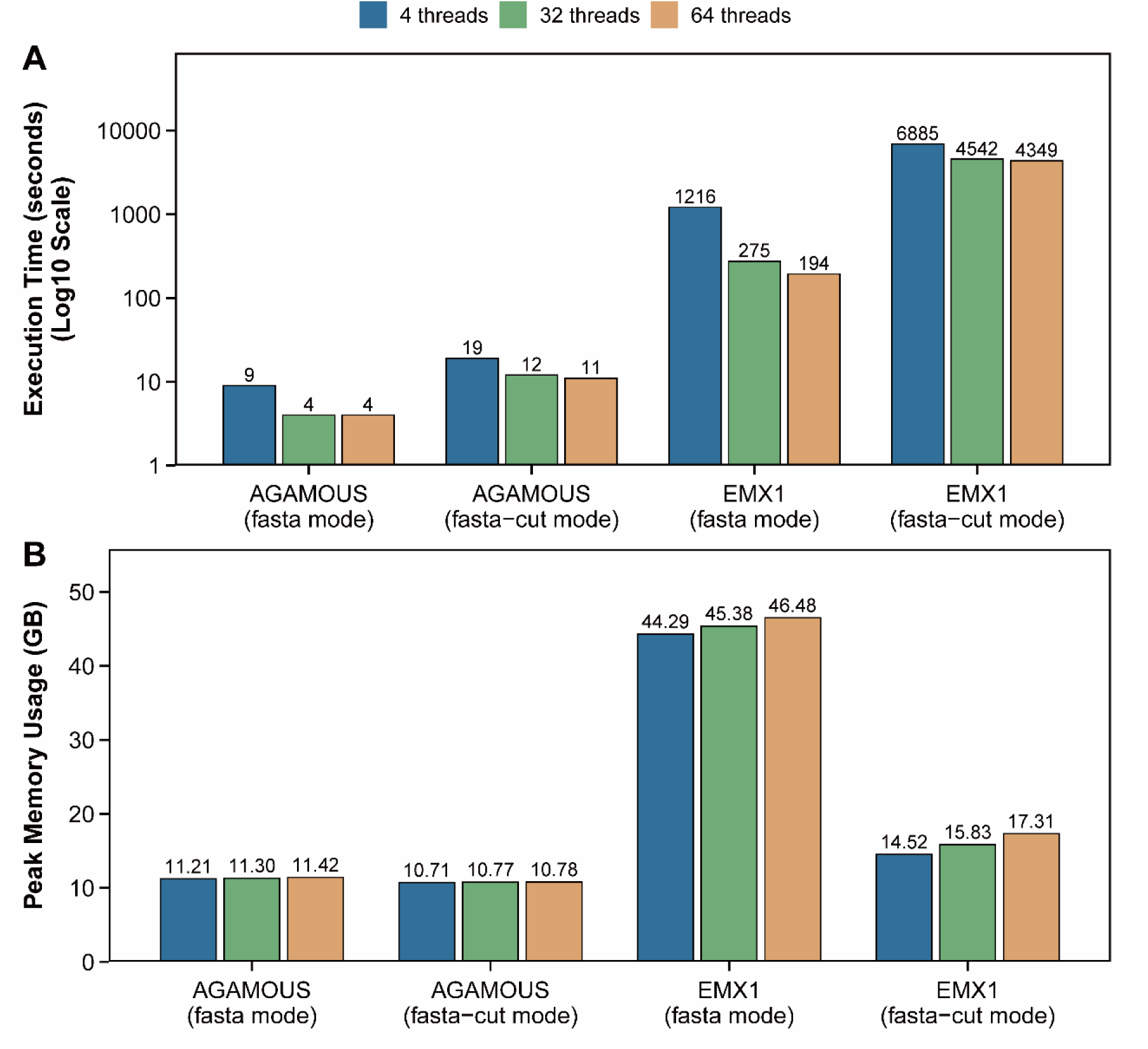
Efficiency of SpacerScope Peak memory usage. Note: The values in the figure represent overall system-wide peak memory usage rather than memory used by SpacerScope alone. No other software was running during testing, and the system’s baseline resident memory usage was approximately 10 GB.

## 4. Discussion

The precision of CRISPR/Cas systems remains the cornerstone of their utility in functional genomics and crop improvement. Existing off-target analysis tools often face a trade-off between computational efficiency and detection sensitivity, and current validation workflows can also be biased by their dependence on predicted candidate sites [7, 8]. SpacerScope was designed to address both of these limitations by combining high-throughput binary vectorization with genome-wide analysis that does not depend on a narrowly preselected candidate set.

### Algorithmic Efficiency via State-Space Compression

A key innovation of SpacerScope is the implementation of binary-channel pre-filtering. Conventional whole-genome off-target search tools such as Cas-OFFinder perform direct sequence comparison under predefined mismatch or bulge constraints [1], and approximate matching generally becomes more expensive as the allowed edit space and candidate volume increase [16]. SpacerScope instead adopts the general logic of bit-parallel approximate matching [17], but adapts it to CRISPR off-target search through binary-channel encoding of nucleotides. This representation reduces the effective state space before downstream alignment and enables aggressive pruning of low-probability loci without sacrificing sensitivity.

### Engineering Optima: Memory Bandwidth vs. Arithmetic Complexity

In developing our substitution-detection module, we evaluated two broad classes of approximate-matching strategies: q-gram-based filtering [18] and bit-parallel matching [17]. Our benchmarks showed that, for short spacers scanned against very large target libraries, the dominant bottleneck is memory bandwidth and cache locality rather than the arithmetic cost of Hamming-distance calculation itself. Under these conditions, batch brute-force matching with compact 2-bit encoding provided the best engineering trade-off: it minimizes per-target memory footprint, preserves sequential access patterns, and avoids the index-construction and reuse assumptions required by q-gram-based methods. We therefore retained 2-bit-encoded batch scanning as the main substitution-detection path in SpacerScope.

### Mitigating Analysis Bias and Indel “Blind Spots”

A significant contribution of this work is the reduction of analysis bias in off-target detection. Plant off-target studies have noted that validation is often restricted to predicted candidate sites rather than unbiased genome-wide analysis [7], which can create a blind spot for true off-target mutations in unmarked genomic regions. In our benchmarks, established tools such as Cas-OFFinder and web-based platforms such as CHOPCHOP showed limited recovery of complex mismatch patterns, including tandem insertions and combined substitution-indel events [1, 2]. SpacerScope addresses this gap with a right-end-anchored edit distance that is motivated by the PAM-proximal asymmetry of CRISPR targeting [19, 20]. By allowing flexible trimming at the PAM-distal side while preserving the PAM-proximal anchor, the method can accommodate length variation patterns that remain biologically relevant in bulge-containing off-targets [21]. We then refine the interpretation of each retained site with an empirical risk-scoring framework that is conceptually aligned with data-driven off-target scoring models [22], yielding an interpretable risk index rather than a raw distance value alone.

### Validation and Future Perspectives

The robustness of this framework is supported by our retrospective analysis of CIRCLE-seq data [11], in which SpacerScope recovered 100% of validated sites within the defined parameter space. The tool also identified additional high-risk sites that were initially misclassified because of more complex edit patterns, highlighting the corrective value of the refined-alignment stage. Computational trade-offs nevertheless remain. When the allowed indel threshold is increased substantially, variant generation grows non-linearly and runtime rises accordingly. In bulge-aware off-target search, practical analyses commonly restrict the search space to one or two bulges [23], and our own rice and CIRCLE-seq benchmarks indicate that an indel threshold of 2 was sufficient under the settings evaluated here. Future iterations could therefore focus on more aggressive pruning heuristics or GPU-accelerated variant generation to expand the searchable parameter space without sacrificing sensitivity.

## 5. Conclusion

SpacerScope establishes a robust, genome-wide off-target profiling workflow that harmonizes bitwise pre-filtering with high-fidelity indel validation. By bridging the gap between high-speed heuristic tools and exhaustive search engines, we provide a versatile solution for large-scale CRISPR analysis in both model organisms and complex polyploid crops. The integration of event-level annotations and empirical risk scoring ensures that SpacerScope is not merely a search engine, but a comprehensive decision-support tool for precision genome engineering.

## 6. Acknowledgment

This work was financially supported by the National Natural Science Foundation of China (32572957). During the code implementation and manuscript preparation for this study, large language models such as ChatGPT were used to assist with code review and language polishing. All technical content, experimental design, data analysis, and conclusions were completed independently by the authors, who take full responsibility for the accuracy and reliability of the final content.

## 7. Author Contributions

Yanji Qu: Software, Validation, Formal analysis, Algorithm development, Writing the manuscript draft. Yaxuan Wang: Validation, Software testing, Review & editing. Yan Wang, Haoru Tang: Software testing, Debugging, reviewing the manuscript. Qing Chen: Conceptualization, Methodology, Supervision, Project administration, Writing – review & editing. All authors have read and approved the final manuscript.

## 8. Data and Code Availability

SpacerScope source code, precompiled executables, and user documentation are available at https://github.com/charlesqu666/SpacerScope.

## 9. Competing Interests

The authors declare no competing interests.

## 10. Ethics Statement

Not applicable.

## Notes

### Competing Interest Statement

The authors have declared no competing interest.

